# Step-specific adaptation and trade-off over the course of an infection by GASP-mutation small colony variants

**DOI:** 10.1101/2020.05.28.122713

**Authors:** Faucher Christian, Mazana Vincent, Kardacz Marion, Parthuisot Nathalie, Ferdy Jean-Baptiste, Duneau David

## Abstract

During an infection, parasites face a succession of challenges, each decisive for disease outcome. The diversity of challenges requires a series of parasite adaptations to successfully multiply and transmit from host to host. Thus, the pathogen genotypes which succeed during one step might be counter-selected in later stages of the infection. Using the bacteria *Xenorhabdus nematophila* and adult *Drosophila melanogaster* as hosts, we showed that such step-specific adaptations, here linked to GASP mutations in the *X. nematophila* master gene regulator *lrp*, exist and can trade-off with each other. We found that nonsense *lrp* mutations had lowered ability to resist the host immune response, while all classes of mutations in *lrp* were associated with a decrease in the ability to proliferate during early infection. We demonstrate that reduced proliferation of *X. nematophila* best explains diminished virulence in this infection model. Finally, decreased proliferation during the first step of infection is accompanied with improved proliferation during late infection, suggesting a trade-off between the adaptations to each step. Step-specific adaptations could play a crucial role in the chronic phase of infections in any diseases that show similar small colony variants (also known as SCV) to *X. nematophila*.

**Importance:** Within-host evolution has been described in many bacterial diseases, and the genetic basis behind the adaptations stimulated a lot of interest. Yet, the studied adaptations are generally focused on antibiotic resistance, rarely on the adaptation to the environment given by the host, and the potential trade-off hindering adaptations to each step of the infection are rarely considered. Those trade-offs are key to understand intra-host evolution, and thus the dynamics of the infection. However, the understanding of these trade-offs supposes a detailed study of host-pathogen interactions at each step of the infection process, with for each step an adapted methodology. Using *Drosophila melanogaster* as host and the bacteria *Xenorhabdus nematophila*, we investigated the bacterial adaptations resulting from GASP mutations known to induce small colony variant (SCV) phenotype positively selected within-the-host over the course of an infection, and the trade-off between step-specific adaptations.

## Introduction

Successful colonization of a host is essential to the lifecycle of pathogens. Over the course of an infection, pathogens face a series of barriers to establishing an infection, each one decisive for the outcome. Depending on the system and on our level of resolution, we can distinguish a variable number of such steps. In the simplest case, parasites need to first encounter and attach to their host, then to overcome the different lines of immune defence to proliferate and establish within the body, and, finally, to transmit to another host (1–3). Each transition between two steps imposes a new challenge that pathogens might overcome with step-specific adaptations. If traits involved in different steps are uncorrelated, independent adaptative mutations may occur and increase in frequency until the pathogen succeeds in each and every step of the infection. But those step-specific adaptations may also trade-off with each other: the pathogenic strains which are successful and dominant at the initiation of the disease might differ from those that are successful and dominant at a given advanced step of an infection. Within-host evolution has been described in many bacterial diseases, such as *Helicobacter pylori* (4), *Staphiloccocus aureus* (5), or the causing agent of melioidosis - *Burkholderia pseudomallei* (6), and the genetic basis behind the adaptations stimulated a lot of interest. However, the studied adaptations are generally focused on antibiotic resistance, rarely on the adaptation to the environment given by the host, and the potential trade-off hindering adaptations to each step of the infection are rarely considered.

Such trade-offs limit our understanding of how pathogens evolve until we precisely describe the way they interact with their hosts at each step of the disease process. The bacteria *Salmonella enterica* is a good example of model where trade-offs have been described (7). The *in vivo* adaptation to a specific host trades off with the transmission to another host (8), and the adaptation to remain in the blood trades off with the ability to infect the gastrointestinal environment by the loss of key gene functions (9). In *Salmonella enterica*, phase variation (i.e. a mechanism for high frequency back-and-forth switch between phenotypes) of the virulence factor impeding fast proliferation over the course of the infection has been selected to avoid this trade-off (10), but this mechanism is not present in all bacterial diseases and it is not clear whether bacterial adaptations for each step are impeded by the trade-off.

The bacterial entomopathogen *Xenorhabdus nematophila* is a tractable model which can help in understanding the consequences of step-specific adaptations on intra-host evolution. In the wild, *X. nematophila* forms a symbiotic association with its vector, the nematode *Steinernema carpocapsae,* which lives in soil and reproduces in insect hosts. Once in an insect gut, the vectors *S. carpocapsae* inject a few cells of their bacterial symbiont into the insect haemolymph. The bacterial population proliferates despite the host immune response, produces toxins to kill it rapidly, and degrades host tissues, which in turn supports nematodes development and reproduction inside the host body cavity. After the insect’s death, the bacteria cannot disperse before the population of nematode vectors grows. At this step of the infection, the bacteria and nematode must share the dead insect as a stock of resources that are no longer replenished; nematodes then often eat the bacteria to ensure their survival (11). Once dispersing nematode offspring are produced, a small number of bacteria associate with them, thanks to a very specific set of three genes, and disperse with the vector (12, 13). Thus, we can recognize at least three discrete steps in the infection (14): one where the bacteria are in a dedicated vesicle of the nematode, waiting for the next infection; one where the bacteria need to survive the insect immune response and destroy host tissue for the establishment of its vector; and a final step where the bacteria need to persist in the decaying insect host to eventually feed and colonise the vector.

For decades the entomopathogen bacteria *Xenorhabdus nematophila* has been described as occurring under two phenotypes resulting from phase variation (15). Bacteria from the phenotype often referred as 1 are mobile/flagellated, produce antibiotics, hemolysines, immune suppressors and toxins, while bacteria from phenotype 2 generally lack these characteristics (16, 17), and are about ten times smaller (18). However, it has recently been shown that those two discrete phenotypes are in fact not due to phenotypic switching but to selection during the infection of mutations in the Leucin-responsive regulatory protein (*lrp*) (18). Even though a variety of *lrp* mutations are known to be selected over the infection, the differences of advantage they could confer is unknown. *lrp* is a strongly conserved global transcriptional regulator widely distributed among prokaryotes and archaea (19–21). In *Escherichia coli*, it is involved in amino acid metabolism, monitors the general nutritional state by sensing the concentrations of leucine and alanine in the cell, and regulates genes involved in entering the stationary phase of growth (22, 23). In fact, *lrp* controls the gene expression of about a third of the genome (24) and acts as a virulence regulator in numerous infectious diseases, including those caused by *Salmonella enterica serovar Typhimurium* (25), *Vibrio cholera* (26, 27), *Mycobacteria* (28, 29), *Clostridium difficile* (30), and *Xenorhabdus nematophila* (31, 32). Comparing *in vitro* phenotypes linked to the presence or absence of the mutation in *lrp* with the cycle of the bacteria, one can make a clear hypothesis on how the mutants are selected over the course of the infection. The characteristics of the strains without mutation in *lrp*, described above, suggest that these bacteria are selected to invade a living host carrying other bacterial competitors that must be eliminated to prepare the environment for the nematode. Consistent with this hypothesis, in nature, nematodes generally carry a clonal bacterial population with phenotype 1, which we below name ‘wildtype strain’ (15). On the other hand, the typical characteristics of bacteria living in an environment limited in resources - stationary phase (33) - correspond to the characteristics of the bacteria in phenotype 2, below named ‘mutant’. This suggests that mutations in *lrp* could give an advantage to wait in a decaying host, where quality and/or quantity of resources or other conditions, such as pH, change. In fact, *lrp* mutants outcompete wildtype in aged *in vitro* cultures of *X. nematophila* (18), as does *Escherichia coli* (34), a phenotype described as growth advantage in stationary phase (GASP) (35). Furthermore, *lrp* mutants can outcompete wildtype *in vivo*, when they colonize mouse gut (36). Thus, a clonal bacterial population selected to initiate the infection could be first favoured until the environment changes, and mutants able to grow in stressful conditions become dominants. These mutants can potentially associate with the nematodes, even though badly, but they have never been found in wild-caught nematodes. However, they may serve as food or process available food to provide nutrients for their vector until the rare wildtype genotype or a genotype with a compensatory mutation, allowing phenotype reversion, can re-associate with the nematodes, disperse from the host cadaver inside the new vector, and initiate a new infection. It is not clear whether acquiring this adaptation to persist in the host would hamper the adaptation to initiate infection.

In this study we investigated the bacterial adaptations resulting from mutations known to be positively selected within-the-host over the course of an infection and the trade-off between step-specific adaptations. More specifically, we characterised the consequences of nonsense and missense mutations in the major regulator gene *lrp* on *Drosophila melanogaster* infection at different steps of the infections. Our results suggest that virulence decreased as mutations in *lrp* become more disruptive to gene function, correlating with the ability to grow at the beginning of the infection. Despite the fact that bacteria carrying nonsense mutations in *lrp* proliferated better in immunodeficient compared to healthy hosts, the ability to cope with immune system activation did not correlate with virulence. Furthermore, mutants killed hosts at similar bacterial loads to wildtype, suggesting that they were as pathogenic as wildtype strains and that the ability to kill was solely explained by the speed of proliferation within the host. Next, we demonstrated that mutants, regardless of the type of mutation, were well-adapted to the waiting step of the infection as they proliferated better in dead hosts than do wildtype. Our results suggest that wildtype strains perform better than mutants during the first step of infection, but less well during the second step. We demonstrated that as the infection progressed, the ability to grow well in dead host was acquired while the ability to grow well in the healthy host was lost, suggesting a trade-off between the adaptations to each step.

## Results

### The bacterium-host model

*Xenorhabdus nematophila* is a generalist, and highly virulent entomopathogen bacterium. It is generally described as having two distinct phenotypes, distinguished in the lab by their capacity to adsorb a dye (bromothymol blue). This phenotype is associated with mutations in the *lrp* gene occurring *in vitro* and *in vivo*, which the wildtype strains do not carry. The strains used in this study were chosen from a collection of 34 strains; some are wildtype for the *lrp* gene (blue color in the following results), while the others carry various classes of mutation in different domains of *lrp* (red in the following results). *D. melanogaster* is not a natural host of our generalist parasite, but the genetic tools this model offers allowed us to test predictions on the role of arthropod host immunity in the success of wildtype and mutant bacteria.

### *lrp* mutations impede virulence

We investigated the difference in virulence between *X. nematophila* strains. Using Canton S wildtypes as host genotype, we found that all strains killed their host in less than two days post-injection, but flies infected with wildtype bacteria died more rapidly than those infected with mutants (Figure 2A; Coxme, χ^2^=21.4, p=3.7e^−06^). Where wildtype bacteria killed on average 50 % of their hosts in approximately 14 hours, mutants killed almost none of their host (~7 %). We computed hazard ratio relative to sham infection (Figure 2B): the risk of death was increased on average by a factor of 10^3.3^ when hosts were infected with wildtype strains, but only by a factor of 102.3 while infected by strains with a missense mutation in *lrp* or 10^1.8^ with strains with a nonsense mutation.

**Figure 1:**
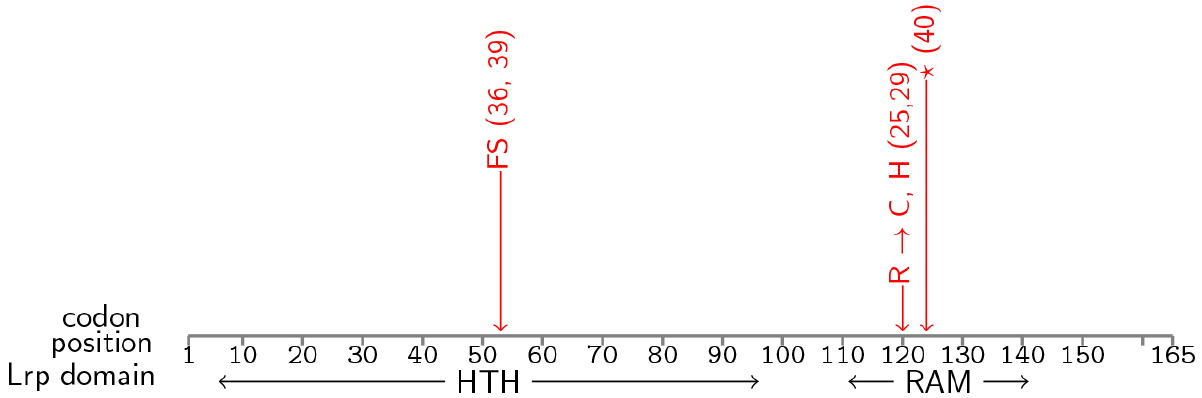
Description of *lrp* mutations. The *lrp* gene is made of two domains, HTH and RAM. The study used four *lrp* mutants each with a single point mutation. Strains 36 and 39 carry different nonsense mutations in codon position 53 leading to a frameshift (FS). Strains 25 and 29 carry each a missense mutation in codon position 120 leading to different amino acids. Strain 40 carries a nonsense mutation (duplication) in codon position 124 leading to a stop codon (*).

**Figure 2:**
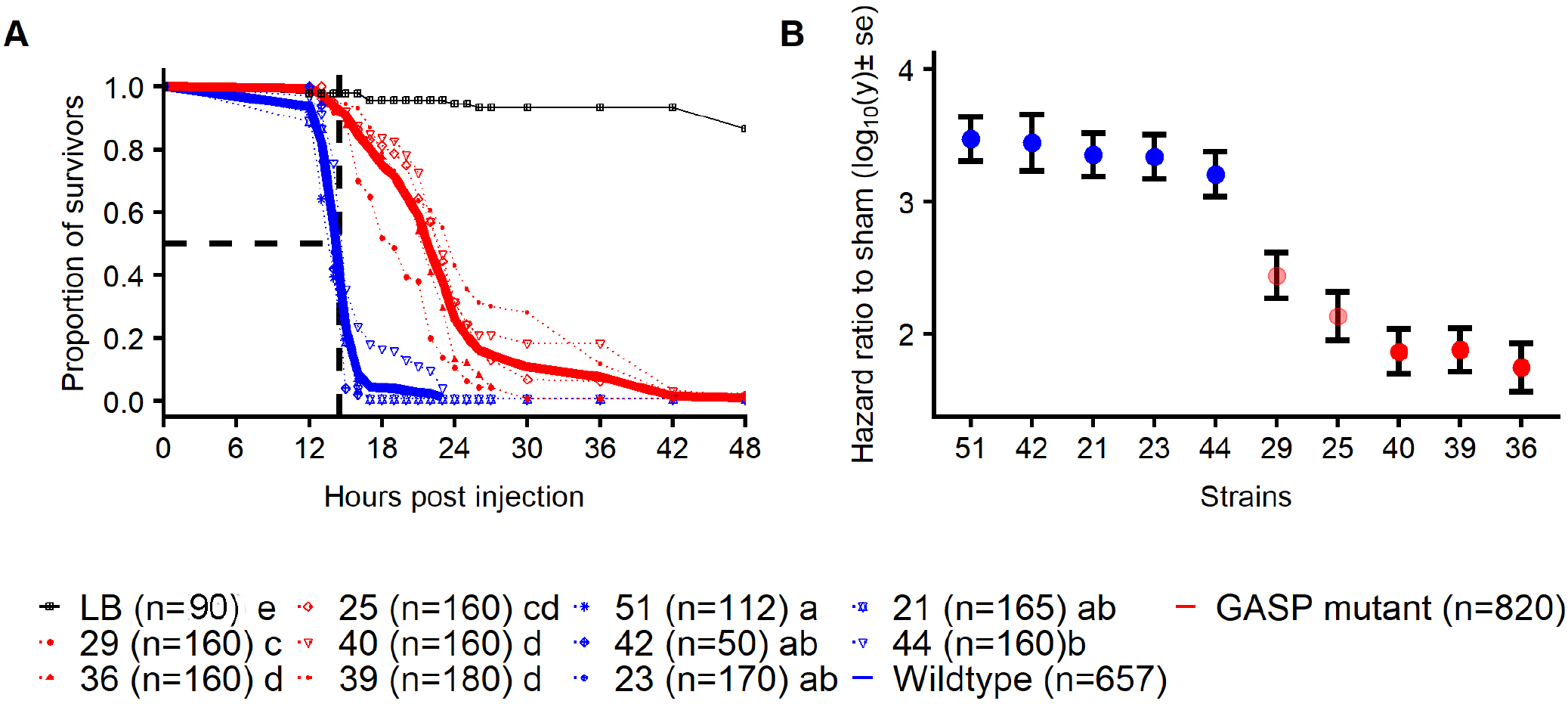
Survival of adult *Drosophila melanogaster* upon infection with wildtype or *lrp* mutant *Xenorhabdus nematophila*. **A-** Survival over time of hosts when injected with 1000 bacteria. Wildtype bacteria, represented individually by blue dotted lines, always killed their hosts faster than *lrp* mutants, represented individually by red dotted lines. Solid lines represent the pooled wildtype (blue) and mutant (red) strains. Black dashed lines show the LT50 of wildtype strains. Number of host individuals are mentioned between brackets. **B-** Virulence of each bacterial strains. Each dot represents the death hazard ratio relative to sham infection calculated from the cox model used to analyse the survival in A. Blue represents infection with wildtype bacterial strains and red the mutant strains. Reference of strain numbers are detailed in Figure 1.

### Wildtype strains grow faster at the beginning of the infection than *lrp* mutants

We hypothesized that if wildtype strains are adapted to initiate the infection, they should grow faster than *lrp* mutants in the early stages of the infection. To test this hypothesis, we compared the bacterial load of hosts infected with four wildtype and four *lrp* mutant *X. nematophila* strains, eight hours post-injection. Overall, hosts infected with wildtype strains had higher bacterial loads 8 hours post-injection (fitme *Mutation* effect: df=1, LRT= 153.39, p-value= 3.13e^−35^, Figure 3A). Hosts infected with wildtype bacterial genotypes had a higher bacterial load at 8 hours post-injection compared to hosts infected with *lrp* mutant strains, the exception being mutant strain 29, bearing a missense mutation, for which the bacterial load was intermediate. After only 8 hours of infection, strains with a nonsense mutation in *lrp* were already at least four divisions behind other strains (Figure 3A). We tested if this difference in bacterial load was linked, at least in part, to the host immune response. As a Gram-negative bacterium, the main immune pathway activated by *X. nematophila* infection in *Drosophila* is the IMD pathway. The IMD pathway is so important in controlling Gram negative infections that individuals lacking the pathway die in few hours from infections that are otherwise benign (37, 38). It is already known that *X. nematophila* triggers a sustainable IMD response upon systemic infection, that immune deficient hosts die earlier, and that hosts immune primed by other Gram-negative bacteria survive longer to subsequent infection by *X. nematophila* (39, 40). Thus, we compared the bacterial load in control healthy hosts to hosts carrying a knock-out mutation in *Dredd*, a critical gene for the activation of the IMD pathway (41), at 8 hours post-injection. We found that bacterial strains with a nonsense *lrp* mutation had a significantly higher bacterial load in immunodeficient hosts than in healthy hosts, while wildtype and missense bacterial strains had the same bacterial load (Figure 3A). To further support the result suggesting that nonsense mutations in the *lrp* gene trigger a susceptibility to the host immune system, to which wildtype strains seem to be fairly resistant, we compared the difference in growth between 0 hour and 8 hours post-injection (Figure 3B). Bacterial proliferation can be represented by the interaction between a difference in bacterial load in time (0 vs 8h) and host genotype (healthy vs immunodeficient). We found that the interaction was strongly significant only for one of the two nonsense mutations (strain 40, Figure 3B). However, even if not significant, we found a trend for a lower proliferation for the nonsense mutation strain 36 and to some extent to the missense mutation strain 29 (Figure 3B). Although not statistically significant, the lower proliferation of the strain 51 (about one division less over 8h) in wildtype hosts was probably due to the fact that this strain proliferates very well in both conditions and even better if the immune system is impeded (Figure 3B). The virulence, expressed as a higher hazard ratio, correlated with the ability to grow at the beginning of the infection (Figure 3C). However, even if nonsense mutations proliferated better in immunodeficient than in healthy hosts (Figure 3D), the ability to proliferate with the immune system, as expressed as the difference in proliferation in healthy and immunodeficient hosts, did not correlate with the virulence (Figure 3E).

**Figure 3:**
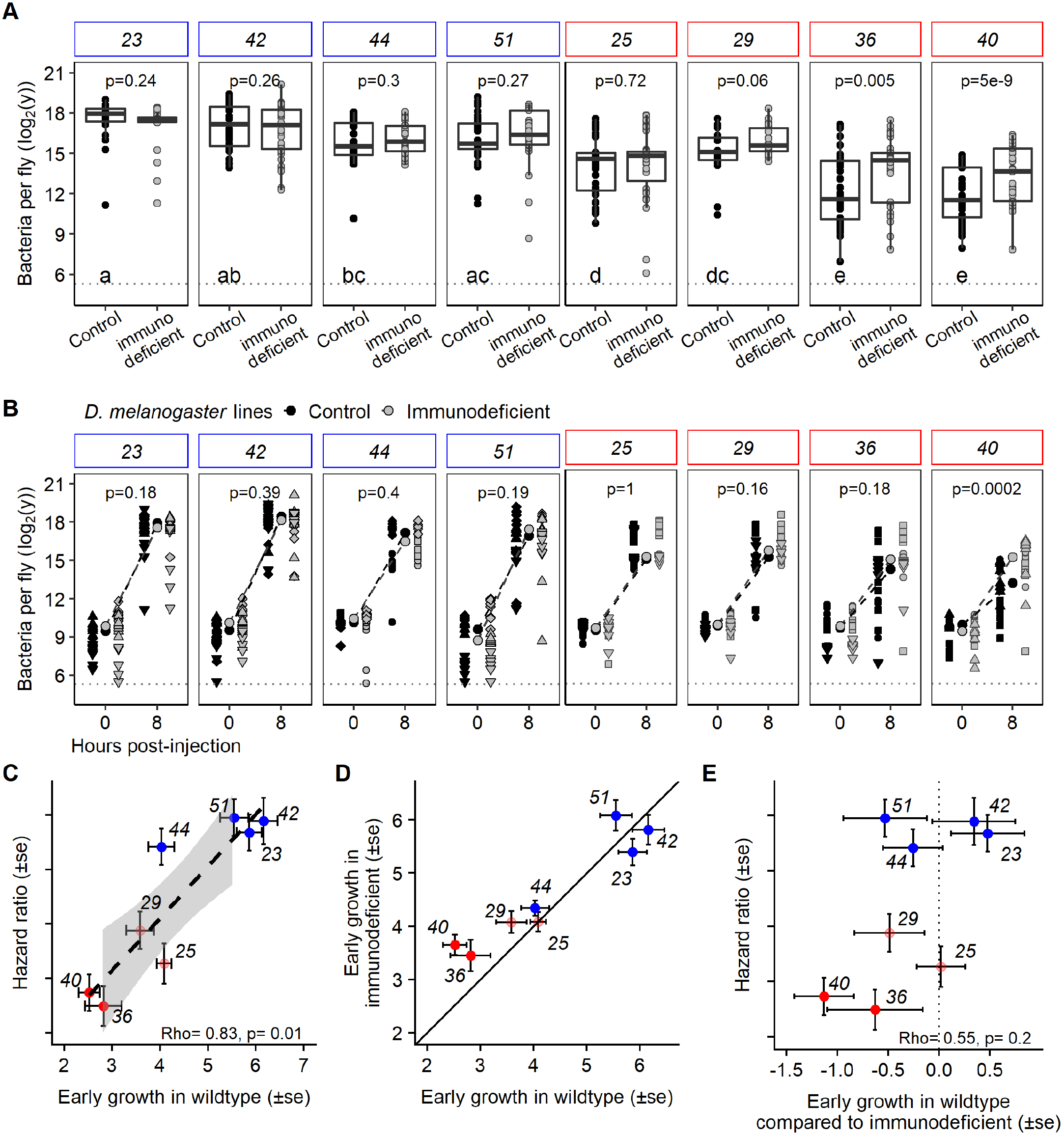
Bacterial success during the first step of the infection. **A-** Bacterial load estimated at 8 hours post-injection in immune (Imd) deficient flies (genotype *Dredd^55^*) and in the genetic background control (*w^1118^*). Letters represent the significant differences between loads of different bacterial strains in hosts with a functional immune system. Wildtype bacterial strains reached a higher density than strains with a mutation in *lrp*. Only mutant strain 29 is intermediate. p indicates the p-value of the difference in bacterial load between immune deficient hosts and healthy hosts. Unlike other strains, those with a nonsense *lrp* mutation proliferated better in immune deficient host. **B-** Bacterial load at injection and 8hours post injection in immune deficient and healthy host. p indicates the p-value of the interaction between time and genotype testing for the difference in proliferation. This approach validates that the strain 40, with a nonsense *lrp* mutation, proliferates less in presence of immune system. **C-** Correlation between early growth in wildtype hosts of the bacterial strains with or without mutation and the hazard ratio (i.e. virulence) extracted for survival analysis in Figure 2. The virulence correlated strongly with the speed of proliferation at the initiation of the infection. **D-** Correlation between early growth in wildtype hosts of the bacterial strains with their early growth in immune deficient host. Solid line represents a perfect correlation (i.e. when the immune system does not affect the bacterial growth). Departure from the line (y=x) indicates a difference in proliferation when the host was immunodeficient. **E-** Correlation between the effect of the immune system on proliferation (i.e. estimate of the interaction effect between time and genotype) and the hazard ratio. Dotted line represents the absence of difference between proliferation in immune deficient hosts and in healthy host. Values are the estimates extracted from the analysis in B. In all panel, blue represents wildtype strains and red represents *lrp* mutants.

### *lrp* mutations do not alter bacterial pathogenicity

The mutations could change virulence via a change in bacterial pathogenicity (i.e. the ability to kill at a given load). We investigated the role of the mutation on pathogenicity by estimating the maximal bacterial load a host can sustain before dying (i.e the Bacterial Load Upon Death or BLUD, see (2)). For a given host genotype, the difference in BLUD between bacterial strains or species reflects the different levels of damage that they inflict to the host, therefore BLUD can be considered a proxy measure of pathogenicity. We hypothesized that the *X. nematophila* wildtype strains, which kill faster, would be more pathogenic (e.g. secrete more toxins) than the mutants and thus, that hosts would succumb to a lower load with wildtype strains than with mutant strains; i.e. the BLUD would be lower in hosts infected by wildtype strains. We found that even if hosts died earlier from infections with wildtype strains than with *lrp* mutant strains, the BLUD was the same (fitme, df=1, LRT = 2.51, p-value=0.11, Figure 4A). As even nonsense mutations had the same BLUD as the wildtype, it is likely that *lrp* does not have a role in the pathogenicity of *Xenorhabdus*. This is further suggested by the absence of correlation between the mean of the hazard ratio of a strain, a proxy for its virulence, and its BLUD mean (Figure 4B) and by previous results showing that *lrp* mutation does not abolish *in vitro* insecticidal secretion activity (40).

**Figure 4:**
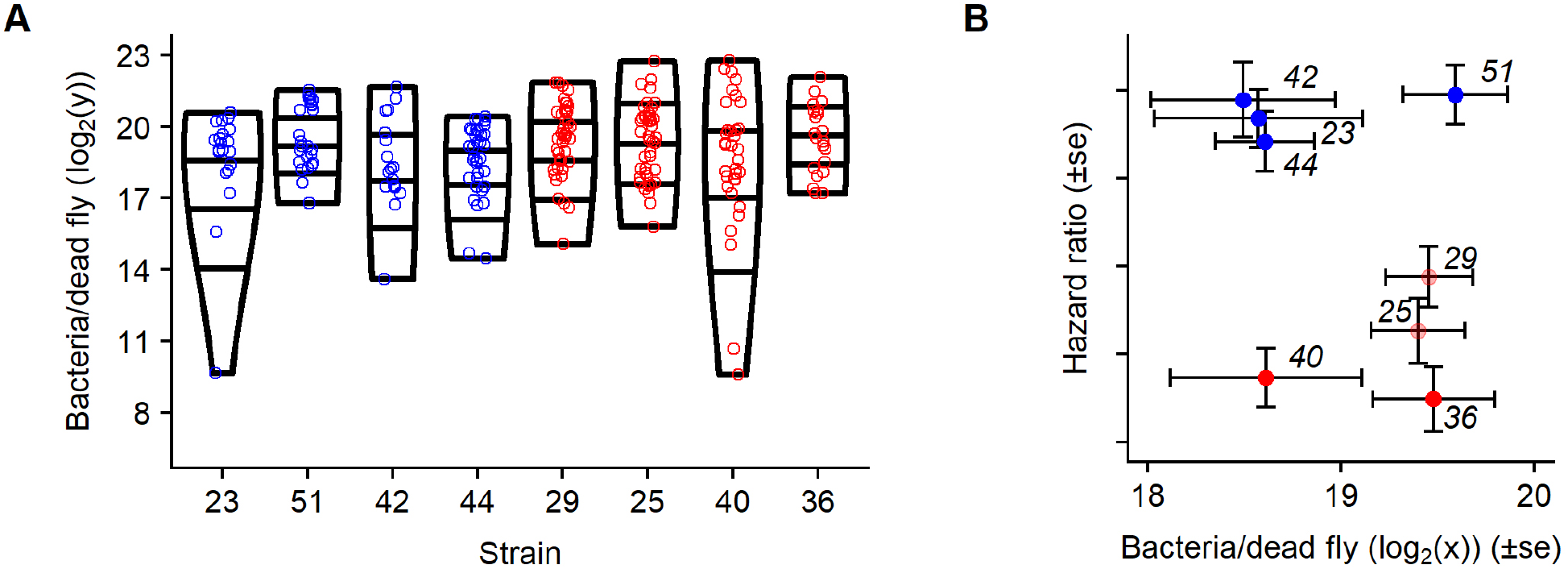
Bacterial load upon death (BLUD) estimated approximately 12h to 30h after injection in wildtype hosts. **A-** Mutant strains reached the same BLUD than wildtype strains. **B-** Correlation between BLUD and hazard ratio. Mutant and wildtype strains reached the same BLUD but wildtype strains had a higher hazard ratio and reached the BLUD about 10 hours earlier, suggesting that *lrp* mutations affect virulence but not pathogenicity.

### *lrp* mutants are better at proliferating in the dead host

We investigated the difference in growth in dead hosts by first estimating the BLUD and then the load 24 hours later. Assuming that *lrp* mutants are selected by the new environment associated with the host death, we hypothesized that they would proliferate better during these 24 hours. All strains were able to grow in dead hosts, however, mutants had a much better ability to grow in those conditions than did wildtype strains (Figure 5A). Interestingly, carrying a missense mutation in *lrp* was sufficient for a strain to be better adapted to this condition, growing at a comparable rate to strains carrying nonsense mutations (Figure 5B).

**Figure 5:**
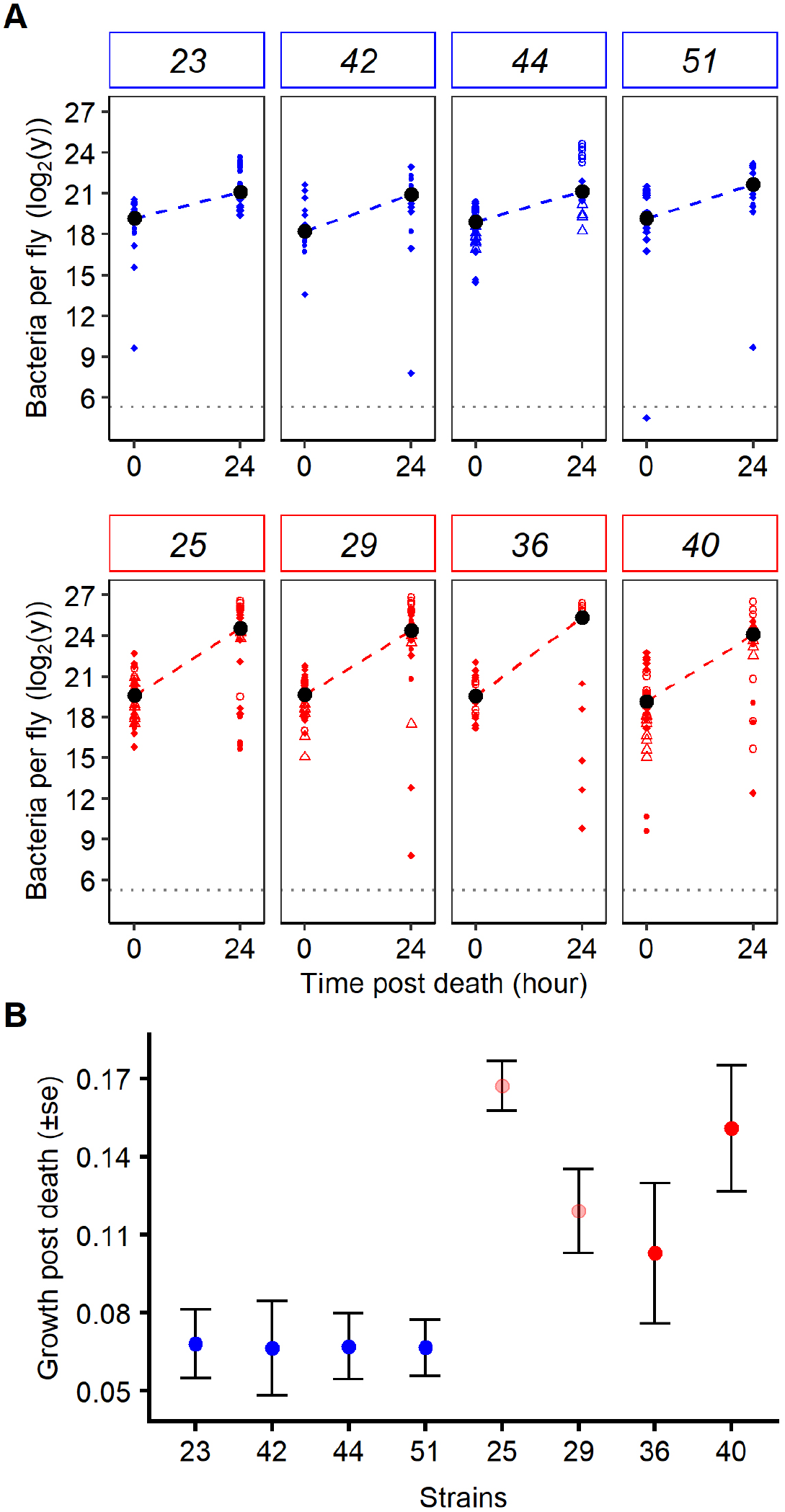
Proliferation during the second step of infection, within the dead host. **A-** Proliferation between death and 24 hours later in wildtype host and **B-** model estimate of growth post death (i.e. estimate of the parameter “Time” in the model to analyse A). Unlike when the host was alive, *lrp* mutant strains had a higher density than wildtype strains when the host was dead. Missense mutation strains (light red) did not proliferate differently in dead hosts than nonsense mutation strains (dark red).

### Trade-off of step-specific adaptations due to *lrp* mutations

There was a clear qualitative trade-off between the ability to proliferate early in the infection and the ability to proliferate in the dead host (Figure 5B & 6). The type of mutation did not determine this trade-off and, even if there was a strong trend, we could not quantitatively show a negative correlation between those two abilities (Figure 6, rho= −0.67, df= 6, t= −2.2, p=0.06). This suggests that changes to, or abrogation of, Lrp function improves performances during the waiting step of the infection but reduces drastically performances at the initiation of the infection.

**Figure 6:**
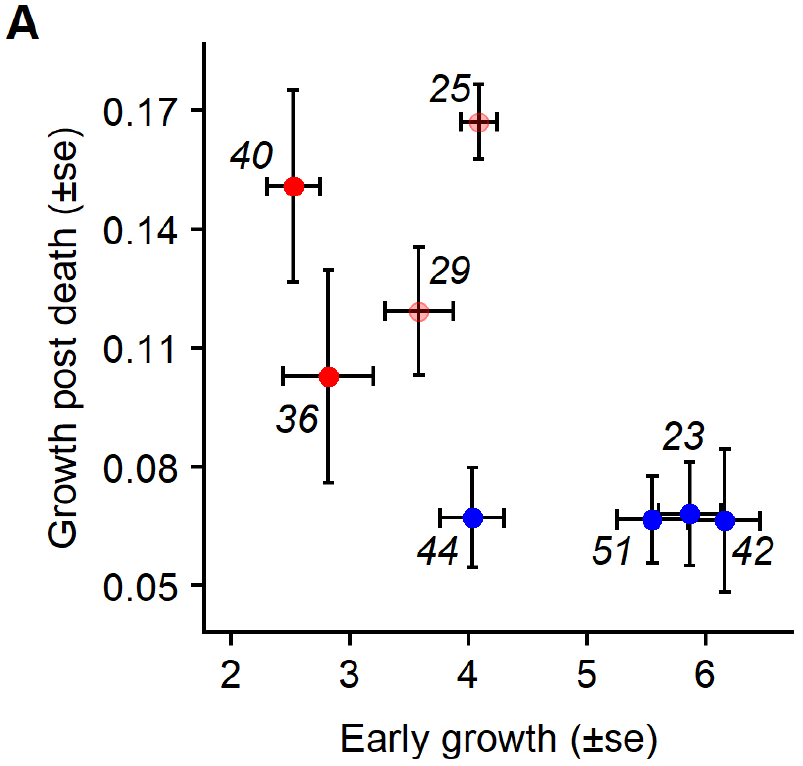
Trade-off between proliferation in first and second step of infection. Correlation between proliferation at the start of the infection and within the dead host. While wildtype strains proliferated the best at the initiation of the infection, they were performing poorly in dead host. Conversely, *lrp* mutant strains proliferated better in dead host but lost the ability to proliferate fast at the initiation of the infection. Correlation between proliferation at the start of the infection and within the dead host. While wildtype strains proliferated the best at the initiation of the infection, they were performing poorly in dead host. Conversely, *lrp* mutant strains proliferated better in dead host but lost the ability to proliferate fast at the initiation of the infection.

## Discussion

At each step of an infection, infected individuals can potentially stop the progress of their parasites. Consequently, there is strong selection on parasites to reach the next step and complete their life cycle (3, 42). However, the traits allowing to progress through each step may be strikingly different and sometimes even be mutually costly. Using *D. melanogaster* as host and *Xenorhabdus nematophila* as a bacterial pathogen, we showed that indeed the bacterial physiological requirements for successive steps of infection are different and can trade-off.

In the system *Xenorhabdus nematophila* (bacteria) – *Steneirnema carpocapseae* (nematode) – insect, the nematode vector injects the bacteria in the insect host body cavity to kill it, and then uses the nutrients of the dead host to reproduce. At the beginning of the infection, bacteria proliferate in a given environment where they need to be highly virulent to prepare the conditions for their vector; once the host is dead, the environment necessarily changes (e.g. in terms of nutrients, oxygen concentration or pH level) and bacteria need to optimize their use of resources that are not replenished until the nematodes produce dispersing larvae. Over the course of an infection, mutations occur in *X. nematophila* populations. The mutations in *lrp* increase in frequency in the population (18), but if they potentially can be carried by the vector, they re-associate poorly (40, 43), possibly because *lrp* affects the expression of genes involved in the mutualism (44). This suggests that wildtype strains, which are those found in wild-caught founder nematodes (15), are adapted to disperse and initiate the infection, while mutants are adapted to persist in the host. We found that both wildtypes and *lrp* mutants proliferated faster *in vivo* at the step they were expected to be better adapted.

The control of bacterial proliferation by the host immune system can take several hours (2) and the acquired immune system has evolved to reduce this crucial time where bacteria proliferate exponentially if untended. Hence, to increase their chance for a successful infection, bacteria can adapt to either suppress, resist, or outpace the humoral immune response (i.e. the control via antimicrobial peptide, AMP, secretion) by killing the host before the AMP concentration is high enough to control its proliferation. Our results show that wildtype strains are most likely better adapted to start the infection because 1- they proliferate faster in the first 8 hours of the infection, 2- they do so at the same speed with or without the host immune response, unlike most *lrp* mutants, and 3- they kill hosts at the same load as *lrp* mutant strains, but do so ten hours earlier. The way in which wildtype bacteria are better adapted to the first step of the infection is still not entirely clear. The efficacy of *Drosophila* cellular immunity in adult is likely to depend on the infection (2, 45) and might not be sufficiently strong to control a virulent pathogen such as *X. nematophila*, unlike the cellular immunity of Lepidopteran. For this reason, our study probably under-estimates the potential adaptation of *X. nematophila* to cellular immunity. Yet, when the *Drosophila* humoral immune response is activated by a pre-exposure to an avirulent bacteria, *X. nematophila* is sensitive to the circulating AMPs (39, 40). This suggests that AMP can control the infection and that *X. nematophila* is not specifically resistant. However, *X. nematophila* is known to not only immunosuppress cellular immunity in Lepidopteran larvae (46, 47) but also downregulate *cecropin*, an AMP (48). Hence, it is likely that wildtype bacteria delay the humoral immune response to kill the host faster. However, *lrp* mutants still kill their host relatively quickly and the ability to grow despite the immune response did not correlate quantitatively with death hazard ratio, which suggests that the efficiency to immunosuppress the host is not the main reason for the higher virulence of the wildtype. However, the death hazard ratio correlated with the speed of proliferation during the first step of infection, which suggest that the adaptation to this step is mainly the intrinsic proliferation rate in this environment.

Although the mutants proliferate in larger numbers in dead hosts showing that they are better adapted to the second step of infection, they re-associate poorly with the nematode (40) and in addition, this adaptation traded-off with the ability to grow at the first step of the infection. If they are not found in wild-caught nematode and are not good at initiating an infection, it is not trivial to understand how mutants are so prevalent in the system. The *lrp* mutants could be selected inside the body of their host, during the infection, with not advantage on the long run, over the whole transmission cycle. This phenomenon is reminiscent of the short-sighted selection of cancerous cells within their host. However, because wildtypes with mutant offspring would have a lower reproductive success, it is likely that any mechanisms preventing those mutations would be advantageous. Alternatively, the occurrence of the mutants may still be important in the symbiosis if they give an advantage to their kin wildtypes. Indeed, *lrp* mutants are likely to modify the nutritional value provided by a dead host and, because they can grow in higher numbers in the dead host, they can feed the vector by being preys. Hence, by feeding the vectors directly or indirectly, the mutants may sustain the system and allow the nematode to disperse with a kin wildtype strain or with a mutant which would have reverted its phenotype.

In our system, the step transition between persistence in a living host and in a dead host is an illustration of the different transitions which can happen in many other diseases, and how trade-off between step-specific adaptations can occur. The change occurring between the acute phase of an infection and its chronic phase is similarly one important and common transition over the course of an infection. One of the bacterial adaptations to this transition is known as small-colony variants (SCV). SCV is important for the chronic establishment of many human diseases such as for instance *Staphilococcus aureus*, *Pseudomonas aeruginosa*, *Salmonella serovar*, *Vibrio cholerae*, *Brucella melitensis*, *Shigella spp.* or *Neisseria gonorrhoeae* (reviewed in Proctor et al., 2006). The pathogen *Staphilococcus aureus* is a good example to illustrate the selection intra-host of SCV as it is its ability to change rapidly from an extracellular aggressive state to SCV adapted for intracellular infection, which allows for long-term persistence in its host. The typical *S. aureus* SCV are typically form small colonies on agar plates, have a decreased respiration, decreased pigmentation, decreased hemolytic activity, decreased coagulase activity, increased resistance to aminoglycoside, and unstable colony phenotype (49). *In vivo*, SCV usually gain in fitness by acquiring the ability to establish and remain chronic in the host after they went through the acute phase of the infection. The phenotype of *X. nematophila* selected during the waiting step is very similar to the commonly described SCV (16, 18). In fact, SCV was observed in *X. nematophila* but not identified as such (50) and the term has been used to describe the alternative phenotype of a closely related species*, Photorhabdus luminescens,* which has the same infection strategy (51–53). The biochemical basis of SCV in mammalian pathogens is generally a single or multiple auxotrophism caused by a simple genetic alteration, exemplified by mutations of genes involved in the biosynthesis of thiamine, menadione, hemin or thymidine (54). Mutations in *lrp* have indeed been found to be involved with induced auxotrophy in *E. coli* and *Komagataeibacter europaeus* (55, 56) and with SCV phenotype in *Pseudomonas aeruginosa* (57). Likewise, mutation in *lrp* is sufficient to switch to SCV-like phenotypes in *X. nematophila*, allowing it to persist in the dead host until the vector disperses. Hence, we believe that our bacteria-insect model system allowed us to study the advantage that SCV can provide over the course of an infection and the trade-off between adaptations of infection steps. As such, it suggests that bacteria initiating a disease can be drastically different from the bacteria selected within the host at later stages of infection, and that intra-host selection is a factor to take into account to understand a pathogen and its weakness during an infection.

## Materials and methods

### Fly stocks and husbandry

*Drosophila melanogaster* were reared on flour-yeast medium (per liter of water: 70 g of corn flour, 70 g of yeast, 15 g of agar, 10 g of vitamins (Vanderzant vitamin mixture for insects from Sigma-Aldrich), 10 g of tegosept -diluted in 20 ml of ethanol- and 3 g of propionic acid). Males between three and nine days old (age was standardized within each experiment) were used in all experiments. Rearing and experiments were conducted at 25°C (±1°C) with a 12 h/12 h light/dark cycle. We used Canton S flies and *w^1118^* inbred, laboratory strains as wildtype genotypes. To test for the susceptibility of the bacteria to the immune system, we used an immune-deficient host line in which Imd pathway signalling is impaired due to a mutation in the gene *Dredd^D55^* (58). *Dredd^D55^* flies, backcrossed in the *w^1118^* genotype, produce lower levels of antimicrobial peptides in response to infection.

### The bacterium

*Xenorhabdus nematophila* is generally described as having two distinct phenotypes, distinguished by their capacity to adsorb bromothymol blue. When plated on a bromothymol agar (NBTA) nutrient (15 g of Nutrient agar, 3 g of beef extract, 5 g of peptone, 8 g of NaCL, 0.04 g of Triphenyl 2,3,5 tetrazolium chloride (TTC) and 25 mg of bromothymol blue per liter of water) wildtype bacteria form blue colonies while others form red colonies. This phenotype is associated in mutations in the *lrp* gene, which the wildtype strains do not carry (16, 18, 32).

Our strains were chosen from a collection of 34 strains isolated from independent *in vitro* culture after several days of a prolongated non-agitated growth in LB medium (Luria Broth, Miller). Although this well characterized collection has been obtained from *in vitro* culture, similar mutants have been found *in vivo* (18). Strains 21, 23, 42, 44, 51 are wildtype for the gene *lrp*. *Lrp* is composed of two domains; a DNA-binding domain called HTH, and a ligand-binding domain called RAM (59) The latter generally interacts directly with amino acids while the former, in response to amino acid concentration, interacts with DNA to modify the expression of hundreds of genes (19, 20). The strains 25 and 29 have a missense mutation (i.e. a single base pair change in the domain RAM of *lrp*, both in codon position 120 - in SNP 358 and 359, respectively - leading to amino acid substitutions). Strains 36, 39 and 40 have an indel mutation in *lrp*. The first two have a frame shift in the HTH domain (in codon position 53), while 40 has a nonsense mutation (i.e. 26 bp insertion in SNP 372 leading to a stop codon) in the RAM domain. Hence, while mutations in strains 25 and 29 are expected to affect the function of the protein, mutations in the strains 36, 39 and 40 are expected to stop its function completely (Figure 1). To summarise, we used two strains of *X. nematophila* per mutation type, such that our results will not be particular of one genotype.

### Quantification of bacterial suspension

*X. nematophila* was grown in liquid LB medium. Overnight cultures, started from a single bacterial colony, were grown to saturation at 28°C in agitated conditions (180 rpm). To prepare the suspension used for injection, we first centrifuged the culture (10000 rpm for 5 min), discarded the supernatant and re-suspended the bacterial pellet in 1 ml of LB, measured optical density (OD_600_) by spectrophotometer, and adjusted by dilution to an OD_600_ of 0.1. To standardize the quantities of bacteria injected inside the host, and because wildtypes and mutants differ in their absorbance, we also used a Thoma cell counting chamber to enumerate cells under a microscope (Olympus BX 51, magnification x200).

### *Xenorhabdus nematophila* injection in *Drosophila melanogaster*

We injected approximately 1000 bacteria per fly in 23 nl of LB medium, between the two first abdominal segments using a Nanoject 2 injection system (Drummond) (60). Controls were injected with 23 nl of sterile LB medium. Prior to injection, flies were anesthetized with CO2. Anesthesia lasted for about five minutes, and flies were observed afterward to confirm they recovered from the procedure.

### Host survival

Upon injection of approximatively 1000 bacteria, flies were kept at 25°C with 60 % humidity, and with *ad libitum* access to food. Survival was scored hourly, starting around 10h post-injection. The dose of bacteria injected was chosen with the rationale that it was within a realistic range for initial load with respect to natural infections, but high enough to be sure that each host was exposed to the bacteria, as very low doses are more prone to random variation in bacterial number, with in some cases no bacteria being injected. All flies died rapidly upon injection, showing that they were all exposed.

### Estimation of bacterial load

To characterize the bacterial within-host dynamics, we quantified the bacterial load in individual flies at two time points during the infection. To extract bacteria from the host, we homogenized individual flies at 30 hertz for 1 minute using a tissue lyser (Tissue Lyser II) in 250 μl of LB medium containing two 2 mm glass beads. At least eight independent replicate measurements (*i.e.* separate extractions on 8 individual flies) were performed per time point, treatment, and experiment. Samples were serially diluted 1 to 1:2500 in 96-well microplates. We then deposited 5 μl drops from each well onto a NBTA plate using a 96 micropipette Integra Viaflo 96. Plates were incubated for 48 hours at 28°C and we then counted the number of colonies that grew within each drop. Such raw data where used in the analysis, but for graphical display we rather used estimations of bacterial loads per fly obtained by adjusting a Poisson generalized linear model where the number of CFU is predicted from dilution. To estimate the Bacterial Load Upon Death (BLUD) (2), infected hosts were checked every 30 minutes and newly dead flies were collected and homogenized, with bacterial load quantified as described above (2). Bacterial load 24 hours after the host death was quantified from individuals kept in closed sterile microtubes.

### Statistical analyses

We carried out all analyses using R (61). We analysed differences in survival (time-to-death curves) using the packages “Survival” and “coxme” (62). We used the *coxme* routine, which allows inclusion of random effects in a Cox model, to model variability among day of experiments and replicated vials. We determined how host survival is affected by bacterial mutations by comparing a Cox model including mutation status as a fixed effect to a model without this fixed effect. We extracted hazard ratio from this survival model to compare survival among strains, taking into account experimental replications. We analysed the differences in bacterial load within the host using general linear mixed models (GLMM) implemented in the package *spaMM* with the function “fitme” (63). We tested the effect of mutation on bacterial load within the host by comparing raw data (*i.e.* the counts of bacterial colony (CFU) in a 5μl drop over several dilutions - only in drops containing less than 50 CFU). Dilution and volume were included in the model as fixed offsets. As before, we modelled fluctuations among experiments or replicate variants as random effects. An additional random effect was added to take into account the count uncertainty among drops for a same individual host.

## Acknowledgments

We thank Sylvie Pagès and Alain Givaudan for the bacterial strains. Luis Teixeira (Instituto Gulbenkian de Ciência, Portugal) for providing *w^1118^* and *Dredd^D55^* host genotypes. Adam Dobson and Jennifer Regan for their comments on the manuscript. DD was supported by the French Laboratory of Excellence project “TULIP” (ANR-10-LABX-41; ANR-11-IDEX-0002-02), by the People Programme (Marie Curie Actions) of the European Union’s Seventh Framework Programme (FP7/2007-2013) under REA grant agreement n. PCOFUND-GA-2013-609102, through the PRESTIGE programme coordinated by Campus France and by the The LIA BEEG-B (Laboratoire International Associé-Bioinformatics, Ecology, Evolution, Genomics and Behaviour) (CNRS). VM was supported by “Investissement d’Avenir” grants managed by the French Agence Nationale de la Recherche (CEBA, ref. ANR-10-LABX-25-01). JBF and NP were supported by “project new Frontiers” from the French Laboratory of Excellence project “TULIP” (ANR-10-LABX-41; ANR-11-IDEX-0002-02).

